# Comparative Insilco functional, structural and phylogenetic characterization of plant Mn-Superoxide Dismutase

**DOI:** 10.1101/2022.05.26.493555

**Authors:** Rashmi Gangwar

## Abstract

The environmental stress is jeopardizing the production as well as the survival of plants. This stress leads to the emergence of ROS, a toxic byproduct of several metabolic processes. To combat the ROS plant possess definite defense machinery, superoxide dismutase is one of the antioxidant enzymes responsible for effectively eradicating these ROS. Amongst several SOD, MnSOD is specialized in eliminating the ROS from mitochondria, a site for cellular energy production. In this study, the MnSOD from 16 plants was considered, and its physiochemical and structural properties were predicted using various computational tools. The physiochemical characterization reveals protein properties such as Mw, PI, AI, GRAVY, and II. The secondary structure and the motif analysis revealed these proteins’ functional characteristics. The 3D structure of several proteins was not available, so the SWISS model was used for the homology modeling. The quality of the predicted protein structures was verified using the Ramachandran Plot, VERIFY 3D, and PROCHECK under the SAVES server. These data would provide a foundation for structural and functional characterization of potential MnSOD yet to be characterized, which is expected to enhance the knowledge about its metal binding and catalytic efficiency for future applications.

## Introduction

Plants thoroughly consume oxygen during various metabolic pathways for the production of energy. There is an escape of individual oxygen molecules during these pathways, which contributes to the generation of highly reactive oxygen species [1, 2]. These ROS are singlet oxygen (1O2), superoxide ions (O22), and peroxides. To a certain extent, these ROS are beneficial to the cell, but beyond a limit, they disturb the redox homeostasis by altering DNA and causing protein oxidation and lipid peroxidation. To overcome the oxidative stress caused by ROS, the plant produces a series of enzymatic and non-enzymatic antioxidants [3]. Superoxide dismutase, an antioxidant enzyme, is considered the first line of defense to scavenge the ROS by converting the singlet oxygen molecule to harmless molecules of hydrogen peroxide, which is later catalyzed into water. These SOD have been categorized into three classes based on different metal ions present at their active center, namely Cu-Zn SOD, MnSOD, and Fe-SOD. Amongst them, MnSOD is of great interest as it is localized in mitochondria, which is a major site for energy metabolism for plant cells [4].

The physicochemical and structural properties of the protein were elucidated through computational tools. Various tools from different sources are available for a vast range of computational analyses to identify and predict a particular protein. The traditional experimental method for characterizing proteins from various organisms has an inevitable drawback: time-consuming, high cost, and requires multiple machines and techniques. Hence these issues can be tackled through Insilco approaches. Most of the necessary information for the characterization and determining the functional and physicochemical properties are attained through amino acid sequences. This study reported the insilico analysis functional expression studies on MnSOD antioxidant proteins of several plant species. To get further insights into the structural features, the modeling was performed that would provide a suggestive idea for its molecular functions.

## Materials and methods

### 2.1 Sequence retrieval from NCBI database

The corresponding amino acid sequences of different manganese superoxide dismutase from 16 different plant species were retrieved from the NCBI (National Centre for Biotechnology Information) database (https://www.ncbi.nlm.nih.gov/) in the FASTA format for all the computational analysis. Only full-length sequences were considered for in silico analysis. The full-length sequence of *Nerium Oleander* was identified in our lab and used for the studies.

### 2.2 Physiochemical characterization

The amino acids are considered to provide an insight into the physicochemical properties of the protein, which can be characterized to evaluate the molecular weight (Expasy compute Mw/PI), theoretical isoelectric point (pI), its extinction coefficient [5], instability index (II – stability of proteins) [6], aliphatic index (AI-relative volume of protein occupied by aliphatic side chains) [7] and grand average hydropathy[8] which were computed using the Expasy’s ProtParam server (http://us.expasy.org/tools/protparam.html).

### 2.3 Secondary structure prediction and motif analysis

The SOPMA (Self Optimized Prediction Method with Alignment) was used for the secondary structure prediction of all the MnSOD proteins, which suitably predicted 69.5% of amino acids for a state description of the secondary structure prediction. The data obtained were curated and submitted in FASTA format for further analysis to ScanProsite (https://prosite.expasy.org/scanprosite/) to analyze the motif present in the protein sequence.

### 2.4 Multiple sequence alignment and phylogenetic tree construction

The multiple sequence alignment of all the protein sequences was done using Clustal Omega software (https://www.ebi.ac.uk/Tools/msa/clustalo/). The obtained FASTA alignment file was used to construct a phylogenetic tree using the Neighbor-joining method using MEGA 10.0 with a bootstrap value of 1000.

### 2.5 Homology modeling and structure prediction

The homology modeling was performed using Swiss Modeler software. The best match structure with the known template was selected and analyzed further. The PDB file generated for each protein structure through modeling was examined using the SAVES server under which the Ramachandran plot, ERRAT plot, and VERIFY 3D were functioned to assess the quality of the predicted protein models.

## 3. Results and discussion

### 3.1 Secondary structure prediction of MnSOD proteins

Antioxidants are a well-known and interesting set of proteins; among them, MnSOD proves to be the primary antioxidant that involves an initial defense mechanism against various biotic and abiotic stress. From the available data of MnSOD proteins in NCBI, 16 were considered for further insilico analysis. One of them is *N.oleander*, which is characterized in our lab. A graphical representation of the percentage of helices, beta-sheet, and random coils is given in Fig 2. The results revealed that overall the percentage of α helices was predominant in each individual MnSOD protein with the highest number in *O.sativa* (54%) and lowest in *S.indicum* (44%), followed by random coil percentage, which was more in *H.annuus* (36.4%), *S.indicum* (34.67%), *C. japonica* (34.21%) the others were in the range of (26-30%). This significantly shows that the MnSOD protein is predominantly an alpha-helical protein irrespective of its difference in origin. The detailed amino acid composition, as shown in Fig 4. The composition of amino acids indicates that the MnSOD protein predominantly consists of non-polar amino acids alanine, leucine, and glycine.

**Figure 1.**
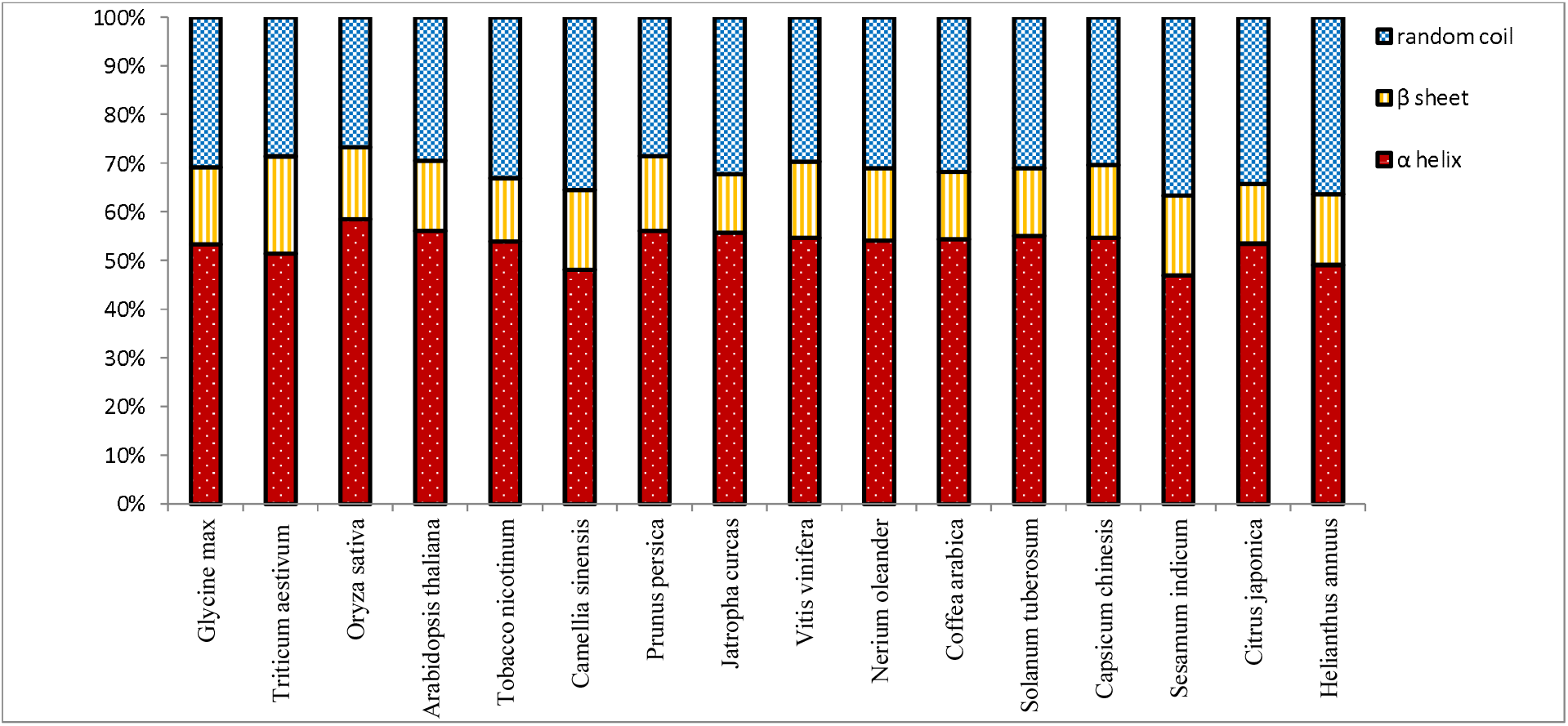
Graphical representation of helixes, sheets, and the random coil of Mn-Superoxide dismutase.

**Figure 2.**
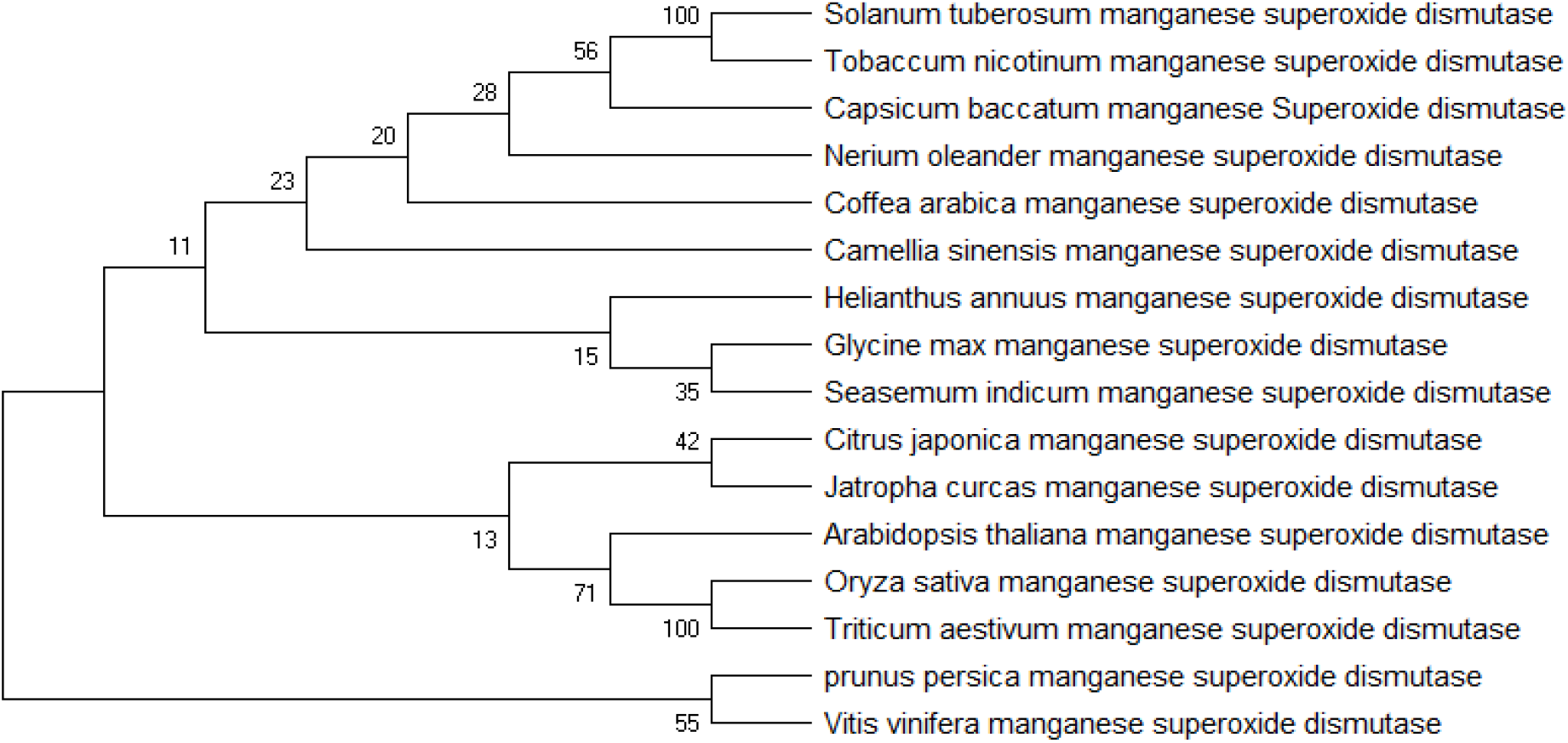
Phylogenetic tree of Mn-Superoxide dismutase proteins was constructed using neighbor-joining method by MEGA X software.

**Figure 3.**
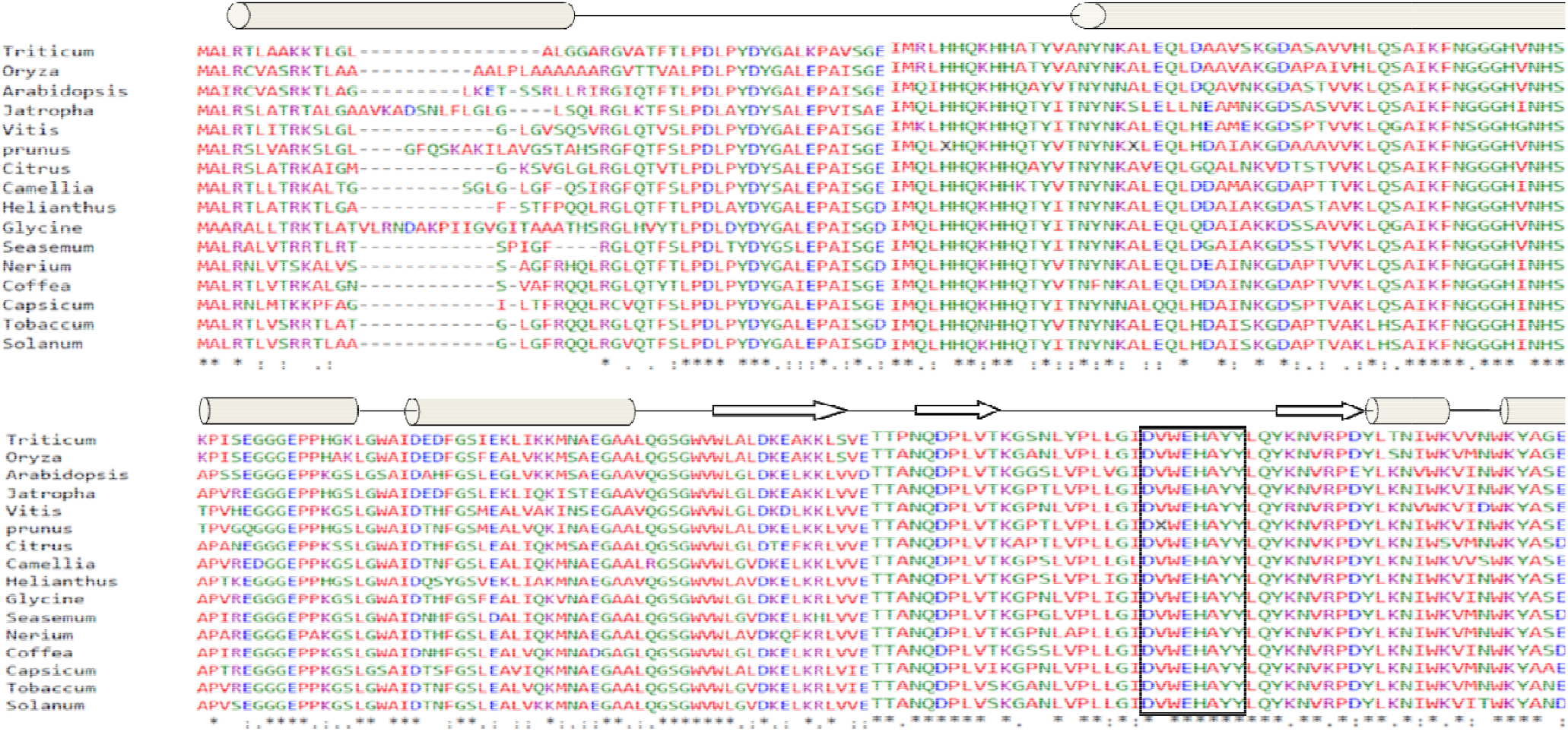
Amino acid sequence alignment of MnSOD. Alpha helices and beta strands are represented as rods and arrows. The conserved MnSOD signature sequence is shown by the box.

**Figure 4.**
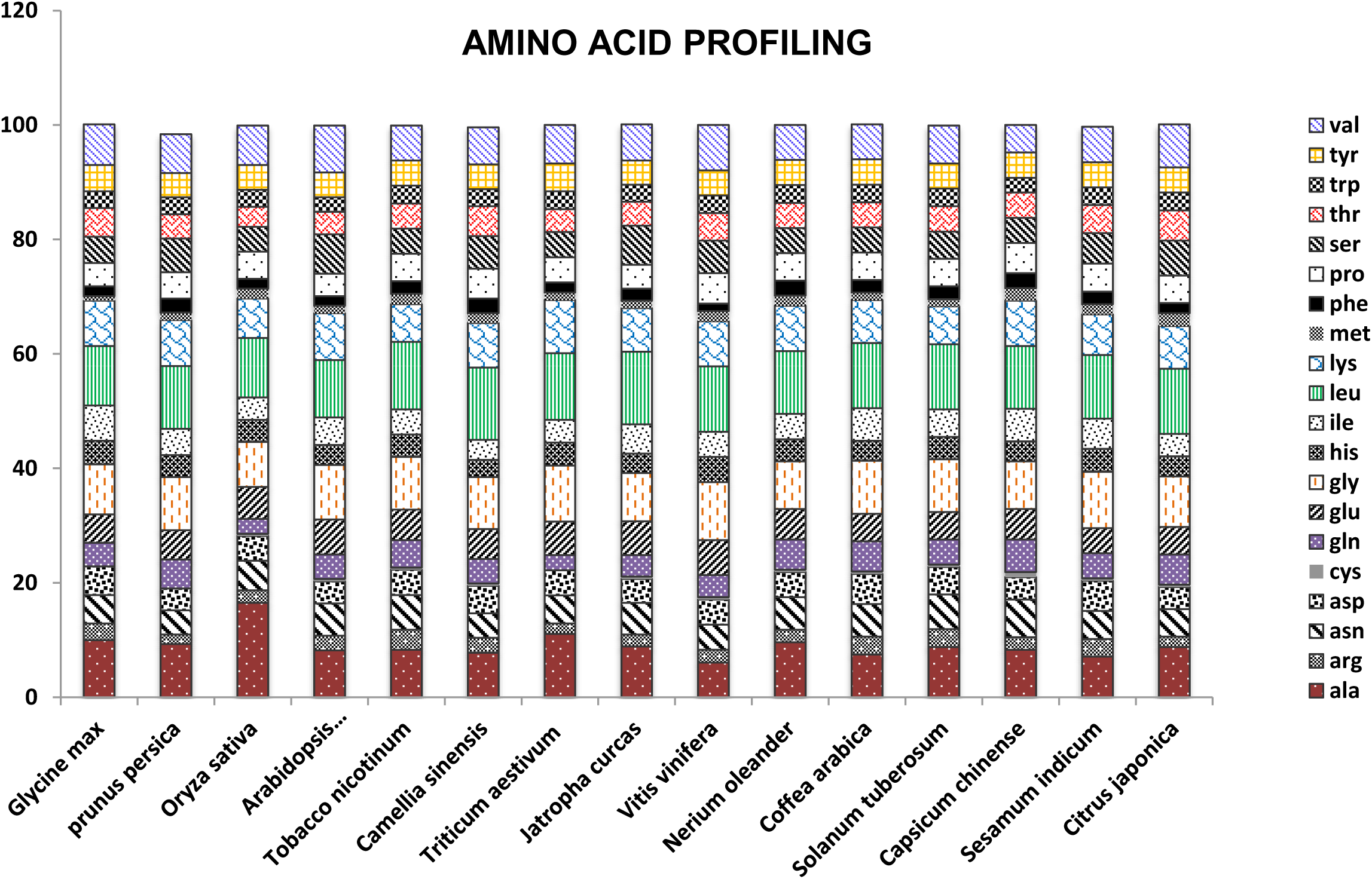
Graphical representation of the percentage of amino acids present in Mn-Superoxide dismutase protein of different plants.

**Figure 5.**
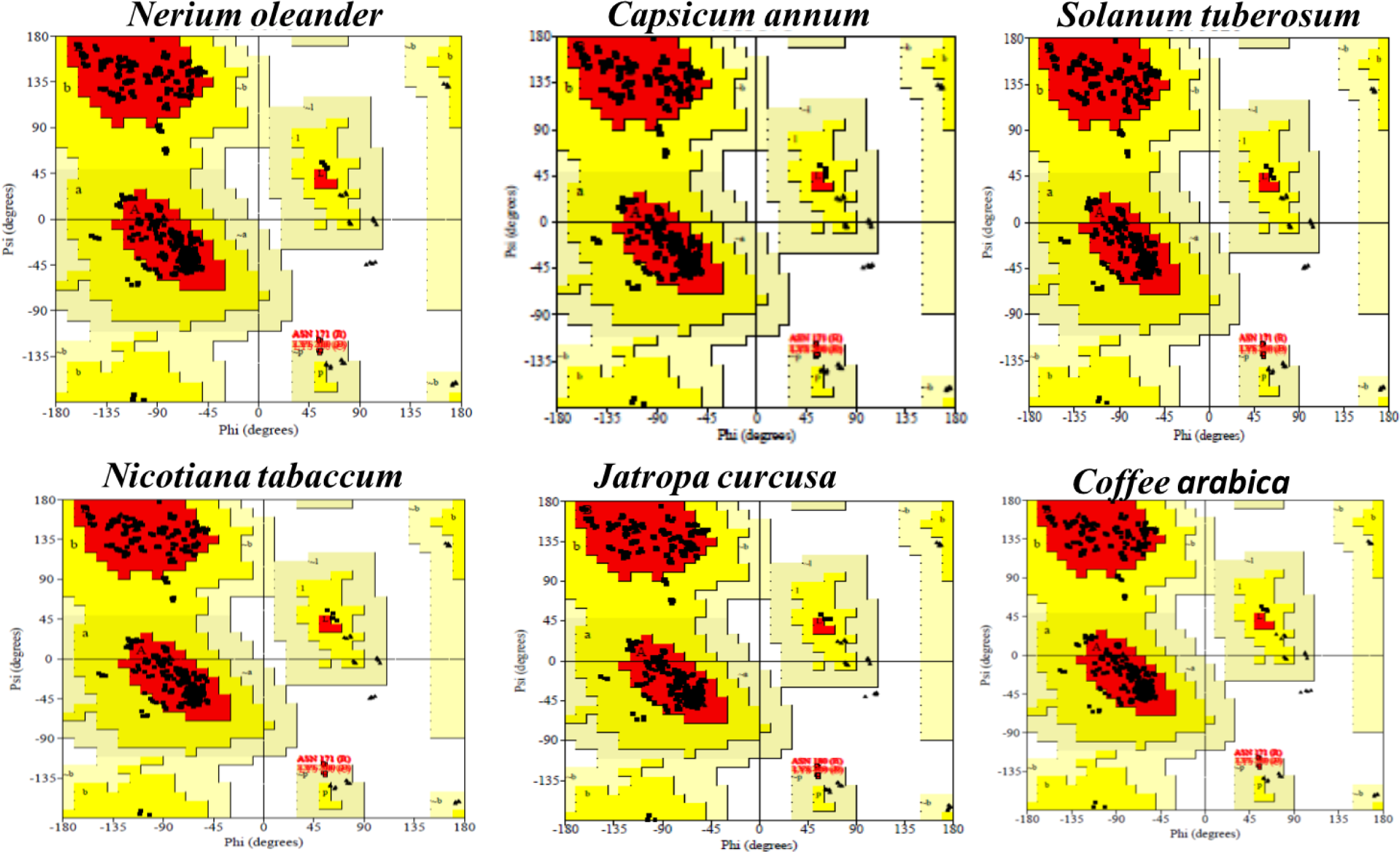
The assessment of the Ramachandran Plot through SAVES server predicting the allowed and disallowed region of several uncharacterized proteins.

### 3.2 Physiochemical characterization

Physiochemical characterization is a crucial aspect of revealing insight into a specific protein. Table 1 shows the different physiochemical properties of all the MnSOD proteins. The total number of amino acid residues lies in the range of 229-241. The *Nerium oleander* also lies in the same range with 228 aa residues, which were found to be similar to *C.arabica, N. tobaccum, S. tuberosum*. The isoelectric point is the condition of a solution where the amino acid produces the same amount of positive and negative charges, resulting in an absolute zero charge on protein. The isoelectric charge of different proteins was above 7, which predicts them to be mildly acidic proteins leaving an exception of *O.sativa* and *V.vinifera* having pI below seven, making them slightly basic. The instability index for all proteins was in a range of 27 to 38, which was below 40. In contrast, the value was 42 in *C.chinesis*, which symbolize all other proteins which have an aliphatic index below 40 were stable, and the one above 40 could be unstable [9]. All proteins showed a higher aliphatic index, which suggested that the protein is possibly thermostable. Here the range of GRAVY laid in between −0.114 and −0.343. Lower the range of GRAVY better would be the interaction between protein and water [10]. The aliphatic index (AI) is defined as the relative volume of a protein occupied by its aliphatic side chains (A, V, I, and L), which is regarded as a confirmatory factor for the increased thermal stability of any globular proteins [7]. The aliphatic index for all the MnSOD protein sequences ranged from 87.32 – to 95.15, which indicates proteins with a high aliphatic index may be thermostable [11]. The extinction coefficient of MnSODs at 280 nm lies within the range of 47900 to 54890 M–1 cm–1 w.r.t. the concentration of Cys, Trp, and Tyr. The high extinction coefficient of *O.sativa, T.aestivum*, and *H.annuus* indicates the presence of a high concentration of Cys, Trp, and Tyr. These computed extinction coefficients of protein help in analyzing the possible protein– protein and protein–ligand interactions in solution.

**Table 1.** Physiochemical properties of Manganese Superoxide dismutase in different plant species.

| **S.No.** | **organism** | **Protein accn. No.** | **aa** | **PI** | **Mw** | **II** | **AI** | **EC** | **GRAVY** | **Half life** |
| --- | --- | --- | --- | --- | --- | --- | --- | --- | --- | --- |
| **1** | *Glycine max* | ABQ52658.1 | 241 | 8.56 | 26.70 | 27.84 | 95.15 | 54890 | -0.285 | 10 |
| **2** | *Triticum aestivum* | AKH40962.1 | 225 | 8.53 | 24.73 | 27.86 | 91.11 | 54890 | -0.285 | 10 |
| **3** | *Oryza sativa* | AAA62657.1 | 231 | 6.50 | 24.99 | 31.67 | 92.25 | 53400 | -0.114 | 10 |
| **4** | *Arabidopsis thaliana* | ACF05703.1 | 231 | 8.47 | 25.44 | 29.27 | 89.48 | 47900 | -0.338 | 10 |
| **5** | *Tobacco nicotinum* | XP_016513256.1 | 228 | 7.86 | 25.50 | 37.16 | 89.43 | 53400 | -0.366 | 10 |
| **6** | *Camellia sinensis* | AJW60327.1 | 230 | 7.81 | 25.56 | 33.46 | 89.48 | 53400 | -0.293 | 10 |
| **7** | *Prunus persica* | XP_007218297.1 | 237 | 8.57 | 26.12 | 29.17 | 89.75 | 53400 | -0.243 | 10 |
| **8** | *Jatropha curcas* | NP_001295624.1 | 237 | 6.71 | 26.21 | 36.10 | 96.33 | 53400 | -0.217 | 10 |
| **9** | *Vitis vinifera* | RVW42331.1 | 228 | 6.80 | 25.31 | 28.67 | 90.61 | 53400 | -0.342 | 10 |
| **10** | *Nerium oleander* | MN365020.1 | 228 | 7.84 | 25.43 | 37.51 | 87.32 | 53400 | -0.352 | 10 |
| **11** | *Coffea arabica* | XP_027110626.1 | 228 | 7.83 | 25.53 | 38.78 | 91.97 | 53400 | -0.363 | 10 |
| **12** | *Solanum tuberosum* | APT43085.1 | 228 | 7.13 | 25.33 | 35.60 | 91.14 | 53400 | -0.313 | 10 |
| **13** | *Capsicum bataccum* | NP_001311927.1 | 228 | 8.39 | 25.57 | 42.28 | 88.60 | 47900 | -0.343 | 10 |
| **14** | *Sesamum indicum* | NP_001306618.1 | 225 | 7.85 | 25.09 | 37.42 | 89.29 | 53400 | -0.336 | 10 |
| **15** | *Citrus japonica* | ADB10839.1 | 228 | 7.83 | 25.15 | 37.82 | 90.26 | 53400 | -0.241 | 10 |
| **16** | *Helianthus annuus* | XP_022006487.1 | 228 | 7.10 | 25.37 | 36.91 | 87.76 | 54890 | -0.339 | 10 |

### 3.3 Sequence alignment and Phylogenetic analysis

The sequence alignment of all the MnSOD proteins reveals that the polypeptide chain is divided into N-terminal helices and a C-terminal α/β domain. All the active sites were located between these N-terminal and C terminal domains with the unique signature sequence pattern D-x-[WF]-E-H-[STA]-[FY] are highly conserved throughout each protein sequence. The phylogenetic tree reveals the distinct relationship amongst each protein in terms of its evolutionary pattern. The whole tree was divided into two major clusters, amongst which the *A.thaliana, T.astevium*, and *O.sativa* belonged to one cluster; the second cluster comprises of all the dicots. The multiple sequence alignment and phylogenetic analysis reveal that the MnSOD sequence possesses high sequence and structural similarity irrespective of their monocot or dicot origin, confirming the common ancestry during Mn-SOD evolution [12, 13]. This evolutionary separation was further observed amongst the dicotyledons and monocotyledons, similarly, as it was found in other MnSOD studies [14, 15]. The protein secondary structure prediction is subjected to a crucial step for full tertiary structure prediction in computational biology.

### 3.4 Fingerprint analysis

The ScanProsite was used for all the fingerprint analysis as it is a vast collection of biologically meaningful signatures that are arranged in the form of patterns which are used for the detection of short motif, or generalized profiles for sensitive detection of larger domains. Each signature is associated with a detailed annotation that provided insights about protein family, domains, or functional sites identified by the signatures. The detailed analysis of all the MnSOD protein sequences revealed that these sequences comprise of following families: PKC_PHOSPHO_SITE (protein kinase phosphorylation sites) that phosphorylates the OH group of serine or threonine and plays a significant role in a wide range of cellular processes like post-translational modification. In plants, protein phosphorylation has been implicated in responses to many signals, including light, pathogen invasion, hormones, temperature stress, and nutrient deprivation. Activities of severa1 plant metabolic and regulatory enzymes are also controlled by reversible phosphorylation[16]. Followed by CK2_PHOSPHO_SITE (casein kinase II (CKII) phosphorylation site), which is a ubiquitous serine-threonine protein kinase highly conserved in all eukaryotes and involved in the regulation of essential cellular processes[17]. N-myristoylation site (N-MYR) leading to the N-terminal myristoylation, which plays a vital role in membrane targeting and signal transduction in plant responses to environmental stress [18]. AMP_PHOPHO_SITE [AMP-activated protein kinase (AMPK)] as AMPK plays critical role in regulating growth and reprogramming metabolism and recently has been connected to cellular processes including autophagy and cell polarity [19]. And lastly, the ASN_GLYCOSYLATION_SITE (N-linked glycosylation) (Table), which is associated with Glycosylation, a significant post-translational modification, and is known to influence protein folding, localization and trafficking, protein solubility, biological activity and cell-cell interactions [20]. In MnSOD, the families PKC_PHOSPHO_SITE, CK 2_PHOSPHO_SITE, N-myristoylation site, AMP_PHOPHO_SITE, and ASN_GLYCOSYLATION_SITE were 2, 4, 6, 4 amino acid in length, amongst them AMP_PHOSPHO_SITE was only present in 4 out of 16 species (*G.max, T.aestivum, O.sativa*, and *A.thaliana*)[21]. The PKC_PHOSPHO_SITE was absent in 3 species and present in rest with a signature sequence of [ST]-x-[RK], variable at 2^nd^ position, and more or less conserved at one and 3^rd^ position. The CK2_PHOSPHO_SITE was found to be absent in *O.sativa* and present in all the remaining species with a signature sequence of [ST]-x (2)-DE]. The four amino acid length sequence was more or less conserved in all the species, but one extra site was present in 7 species, as mentioned clearly in Table 2. The N-myristoylation site (N-MYR) was conserved in all the species throughout with a consensus sequence of G-{EDRKHPFYW}-x (2)-[STAGCN]-{P} in which G is the N-myristoylation site. N-MYR controls the function of the plant protein complex SnRK1, described as one of the most crucial plant regulatory proteins in stress and energy signaling. This lipid modification could be involved in the control of the redox imbalances originating from different stresses in plants [22]. On the other hand, the AMP_PHOSPHO_SITE, which is found in humans, was only present in four species that belong to the dicot plants. The ASN glycosylation with a consensus sequence of Asn-Xaa-Ser/Thr was conserved in the MnSOD belonging to different plant irrespective of its diversity. N-Glycosylation is majorly involved in numerous biological processes which comprises of folding of proteins, protein stability including protein-protein interactions, also the glycan dependent quality assessment processes in the endoplasmic reticulum. Data for A. thaliana and for monocots like rice strongly indicate that complex N-glycans are crucial optimum growth under stressful conditions [23]. The recent identification of rice XYLT-deficient plants with significant growth defects at low temperature is one example [24]. There is an extra glycosylation site present only in *Jatropha curcas* and rest has one with similar pattern. Presence of more than one glycosylation sites leads to structural heterogeneity due to large number of possible stereo- and regio-isomers. This study provides insights about the structural and functional correlation of different plants MnSOD in relation to plant metabolism.

**Table 2.** Fingerprint analysis for MnSOD in plant species.

| **Signature** | **Plant species** | **Sequence** |
| --- | --- | --- |
| PKC_PHOSPHO_SITE | *Glycine max* | TrK |
|  | *Oryza sativa* | SrK |
|  | *Arabidopsis thaliana* | S[r/s]K |
|  | *Tobacco nicotinum* | SrR |
|  | *Camellia sinensis* | [T/S][r/I/V/W][R/K] |
|  | *Vitis vinifera* | [T/S][r/V][K/R] |
|  | *Nerium oleander* | TsK |
|  | *Coffea arabica* | TrK |
|  | *Solanum tuberosum* | [S/T][r/W][R/K] |
|  | *Capsicum bataccum* | T[k/F][K/R] |
|  | *Sesamum indicum* | T[r/I]R |
|  | *Citrus japonica* | TrK |
|  | *Helianthus annuus* | TrK |
| CK2_PHOSPHO_SITE | *Glycine max* | TlpD |
|  | *Triticum aestivum* | [T/S][l/K][p/G]D |
|  | *Arabidopsis thaliana* | [T/S][l/A][p/I]D |
|  | *Tobacco nicotinum* | S[l/K][p/G]D |
|  | *Camellia sinensis* | S[l/A][p/I][D/E] |
|  | *Prunus persica* | SlpD |
|  | *Jatropha curcas* | S[l/A][p/I][D/E] |
|  | *Vitis vinifera* | SlpD |
|  | *Nerium oleander* | TlpD |
|  | *Coffea arabica* | TlpD |
|  | *Solanum tuberosum* | S[l/K][p/G]D |
|  | *Capsicum bataccum* | S[l/A][p/I]D |
|  | *Sesamum indicum* | SlpD |
|  | *Citrus japonica* | [T/S][l/A][p/I][D/E] |
|  | *Helianthus annuus* | TlpD |
| N-myristoylation site | *Glycine max* | G[G/V/I/S][G/T/H/F][I/A/V/E][T/A/N][A/H/L] |
|  | *Triticum aestivum* | G[L/G][A/H][L/R/V][G/N][G/V/H] |
|  | *Oryza sativa* | G[G/S][H/F][V/E][N/A][H/L] |
|  | *Arabidopsis thaliana* | G[L/G/S][K/H/L][E/V/G][T/N/S/G][S/H/A/L] |
|  | *Tobacco nicotinum* | G[G/S][H/L][I/E][N/A][H/L] |
|  | *Camellia sinensis* | G[S/L/G][G/H/L][L/I/E][G/N/A][L/F/H] |
|  | *Prunus persica* | G[G/Q/S][H/G/M][V/G/E][N/G/A][H/E/L] |
|  | *Jatropha curcas* | G[L/G][G/H][L/I][S/N][Q/H] |
|  | *Vitis vinifera* | G[L/V/G/S][G/S/H/M][G/V/Q/G/E][G/S/N/A][V/Q/H/L] |
|  | *Nerium oleander* | G[G/S][E/L/H][P/E/I][A/N][K/L/H] |
|  | *Coffea arabica* | G[N/G/S/L][S/H/L/Q][V/I/A/G][A/N/S][F/H/L/G] |
|  | *Solanum tuberosum* | G[G/S][H/L][I/E][N/A][H/L] |
|  | *Capsicum bataccum* | G[G/S][H/L][I/G/E][N/S/A][H/A/V] |
|  | *Sesamum indicum* | G[G/S][H/L][V/D][N/A][H/L] |
|  | *Citrus japonica* | G[M/Q/G/S][G/A/H/L][K/L/V/E][S/N/A][V/K/H/L] |
|  | *Helianthus annuus* | G[A/G][F/H][S/V][T/N][F/H] |
| AMP_PHOSPHO_SITE | *Glycine max* | KKdS |
|  | *Triticum aestivum* | KKIS |
|  | *Oryza sativa* | KK[m/I]S |
|  | *Arabidopsis thaliana* | KKmS |
| ASN_GLYCOSYLATION | *Glycine max* | NHSI |
|  | *Triticum aestivum* | NHSI |
|  | *Oryza sativa* | NHSI |
|  | *Arabidopsis thaliana* | NHSI |
|  | *Tobacco nicotinum* | NHSI |
|  | *Camellia sinensis* | NHSL |
|  | *Prunus persica* | NHSI |
|  | *Jatropha curcas* | NH/KSI/L |
|  | *Vitis vinifera* | NHSI |
|  | *Nerium oleander* | NHSI |
|  | *Coffea arabica* | NHSI |
|  | *Solanum tuberosum* | NHSI |
|  | *Capsicum bataccum* | NHSV |
|  | *Sesamum indicum* | NHSI |
|  | *Citrus japonica* | NHSI |
|  | *Helianthus annuus* | NHSI |

### 3.5 3D prediction of the MnSOD proteins

The 3D structure provides an insight into the essential characteristics of the protein, so the structural predictions were done for the protein, which lacks such data. The proteins which requires detailed experimental structures were considered. The modeling of the three-dimensional structure of the protein was performed by the Swiss Model, a homology modeling program. The Ramachandran Map generated through the PROCHECK program under the SAVES server with the refinement process assessed the stereochemical quality as well as the accuracy of the predicted protein models. Verify 3D has depicted more than 95% of the residues with an average 3D-1D score greater than 0.2. This demonstrated that the folding energy patterns of the predicted models are in complete agreement and also supported the correctness of the predicted model. The overall favored region, additionally favored and not favored regions of the models were listed in Table 3. The main chain parameters plotted are Ramachandran plot quality, peptide bond planarity, Bad non- bonded interactions, main chain hydrogen bond energy, C-alpha chirality, and over-all G factor. The regions of the quadrangle were considered for the assessment of the residues in the Ramachandran plot analysis. The most allowed regions were shown as red regions in the graph, whereas the additionally allowed regions were visualized as the yellow regions. The Glycine was denoted by triangles whereas the other residues were represented by squares. The distribution of the main chain bond lengths and bond angles were found to be within limits for these proteins. *N. oleander, J. curcusa, N. tabaccum* emerges to have a stable structural moiety in comparison to *Arabidopsis thaliana*, which was considered as the template for building the structure. The more the favored region, the more will be the stability of the predicted model. A good quality model would expect to have over 90% residues in the most favored region (A, B, L) and additional allowed region. These figures which were designated by the Ramachandran Plot, represented them to be good quality predicted models. Furthermore, the ProSA server provides the Zscore value, which lies within the range of scores generally found for the protein native 3D structures, determined by NMR and X-ray crystallography. ProSA gave a Z-score value ranging from −7.26 to −8.43.

**Table 3.** The statistical analysis of the predicted protein through Swiss plot and Ramachandran Plot.

| **S.No.** | **Protein** | **Favored Region (%)** | **Additionally allowed Region (%)** | **Generously Allowed Region (%)** | **Disallowed Region (%)** | **gmqe** | **qmean** |
| --- | --- | --- | --- | --- | --- | --- | --- |
| **1.** | *Coffee arabica* | 92.5 | 6.4 | 0.7 | 0.4 | 0.81 | -0.66 |
| **2.** | *Glycine max* | 92.2 | 6.7 | 0.7 | 0.4 | 0.77 | -0.04 |
| **3.** | *Triticum astevium* | 92.5 | 6.4 | 0.7 | 0.4 | 0.8 | -0.42 |
| **4.** | *Vitis vinefera* | 92.1 | 6.7 | 0.7 | 0.4 | 0.79 | -0.45 |
| **5.** | *Jatropha curcusa* | 92.8 | 6.1 | 0.7 | 0.4 | 0.76 | -0.27 |
| **6.** | *Camellia senensis* | 91.8 | 6.5 | 1.2 | 0.6 | 0.82 | 0.36 |
| **7.** | *Nerium oleander* | 92.7 | 6.1 | 0.6 | 0.6 | 0.81 | 0.14 |
| **8.** | *Prunus persica* | 91.7 | 7.2 | 0.7 | 0.4 | 0.78 | -0.38 |
| **9.** | *Oryza sativa* | 91.9 | 6.7 | 1 | 0.4 | 0.78 | -0.45 |
| **10.** | *Capsicum chinesis* | 92.8 | 6 | 0.7 | 0.4 | 0.81 | -0.07 |
| **11.** | *Solanum tuberosum* | 93.2 | 5.6 | 0.6 | 0.6 | 0.8 | -0.3 |
| **12.** | *Sasemum indicum* | 92.8 | 6.1 | 0.6 | 0.6 | 0.82 | -0.36 |
| **13.** | *Helianthus annus* | 92.4 | 6.4 | 0.7 | 0.4 | 0.8 | -0.39 |
| **14.** | *Citrus japonica* | 91.2 | 6.8 | 1.6 | 0.4 | 0.8 | -1.11 |
| **15.** | *Nicotiana tabaccum* | 93.2 | 5.6 | 0.7 | 0.4 | 0.81 | 0.11 |
| **16.** | *Arabidopsis thaliana* | 91.8 | 7.0 | 0.7 | 0.4 | 0.92 | 0.03 |

## Conclusion

In this study, MnSOD protein of different plant species were selected and characterized to deduce its Physicochemical properties by predicting there expected molecular weight, isoelectric point (pI), EC(extinction coefficient), II(instability index), AI (aliphatic index) and GRAVY (grand average hydropathy). For these proteins, disulphide linkages, motifs, and profiles were predicted. The secondary structure prediction revealed that the alpha helix is predominant in all the proteins followed by random coils. The homology modeling through the Swiss Model gave an insight into the predicted structural aspects of these MnSOD proteins. Therefore the computational analysis provided useful information about the structure and functional aspects of these MnSOD proteins where the data was not available due to the unavailability of the crystal structures.

## Conflict of Interest

The authors share no conflict of interest.

## Acknowledgments

This research was financially supported in the form of fellowship to Ms. Rashmi Gangwar by the Department of Science and Technology (DST) under the INSPIRE scheme, grant no IF140298.

## Notes

### Competing Interest Statement

The authors have declared no competing interest.

### Summary of Updates

The email address of the corresponding author was incorrect.

